# Assessment of contaminants, health and survival of migratory shorebirds in natural versus artificial wetlands – the potential of wastewater treatment plants as alternative habitats

**DOI:** 10.1101/2023.02.15.528752

**Authors:** Tobias A. Ross, Junjie Zhang, Michelle Wille, Tomasz Maciej Ciesielski, Alexandros G. Asimakopoulos, Prescillia Lemesle, Tonje G. Skaalvik, Robyn Atkinson, Roz Jessop, Victorian Wader Study Group, Veerle L. B. Jaspers, Marcel Klaassen

## Abstract

The rapid destruction of natural wetland habitats over past decades has been partially offset by an increase in artificial wetlands. However, these also include wastewater treatment plants, which may pose a pollution risk to the wildlife using them. We studied two long-distance Arctic-breeding migratory shorebird species, curlew sandpiper (*Calidris ferruginea*, n=70) and red-necked stint (*Calidris ruficollis*, n=100), while on their Australian non-breeding grounds using a natural wetland versus an artificial wetland at a wastewater treatment plant (WTP). We compared pollutant exposure (elements and per- and poly-fluoroalkyl substances/PFASs), disease (avian influenza), physiological status (oxidative stress) of the birds at the two locations from 2011-2020, and population survival from 1978-2019. Our results indicated no significant differences in blood pellet pollutant concentrations between the habitats except mercury (WTP median: 224 ng/g, range: 19-873 ng/g; natural wetland: 160 ng/g, 22-998 ng/g) and perfluorooctanesulfonic acid (WTP median: 52 ng/g, range: <0.01-1280 ng/g; natural wetland: 14 ng/g, <0.01-379 ng/g) which were higher at the WTP, and selenium which was lower at the WTP (WTP median: 5000 ng/g, range: 1950-34400 ng/g; natural wetland: 19200 ng/g, 4130-65200 ng/g). We also measured higher blood o,o’-dityrosine (an indicator of protein damage) at the WTP. No significant differences were found for adult survival, but survival of immature birds at the WTP appeared to be lower which could be due to higher dispersal to other wetlands. Interestingly, we found active avian influenza infections were higher in the natural habitat, while seropositivity was higher in the WTP, seemingly not directly related to pollutant exposure. Overall, we found negligible differences in pollutant exposure, health and survival of the shorebirds in the two habitats. Our findings suggest that appropriately managed wastewater treatment wetlands may provide a suitable alternative habitat to these migratory species, curbing the decline of shorebird populations from widespread habitat loss.

## Introduction

Wetlands, both inland and coastal, have experienced one of the most rapid declines of any ecosystem on the planet, with half to two thirds of coastal wetlands having been destroyed in the last 50 years alone (Finlayson & Rea, 1999; Murray, Clemens, Phinn, Possingham, & Fuller, 2014). Rapid human expansion is the major driving force behind these declines, and perhaps no more acutely is this felt than in coastal wetland habitats. In addition to wetlands, more than 16% of tidal flat habitat has been destroyed between 1984-2016 (Murray et al., 2019), though in some areas coastal habitat loss over a similar time period has been shown to exceed 65% (Murray et al., 2014). The effects of widespread wetland destruction are particularly evident in the East Asian-Australasian Flyway (EAAF) (Ma et al., 2014; Murray et al., 2014). Chains of wetlands within this flyway provide vital staging and fuelling areas for migratory shorebirds (Order *Charadriiformes*) during their biannual migrations. Unfortunately, reliance on these sites makes these birds particularly prone to the widespread destruction of wetlands (Lisovski, Gosbell, Minton, & Klaassen, 2020; Murray et al., 2018). As a result, shorebirds are among the most threatened bird families in the world. Indeed, some populations have declined by up to 80% in the last 30 years alone (Conklin, Verkuil, & Smith, 2014; Garnett, Szabo, & Dutson, 2011). Somewhat compensating for the loss of natural wetlands is that the number of artificial wetlands is increasing, including the number of waste stabilisation ponds. Many sewage treatment plants carry out at least some treatment of human waste in open-air settling ponds that provide consistent water availability and high nutrient density. These habitats appear to be of increasing value to a wide range of bird species (Murray, Kasel, Loyn, Hepworth, & Hamilton, 2013), providing a unique opportunity to both manage human waste as well as combat habitat loss for wetland-dependent wildlife.

Potentially compounding the threat of habitat loss is growing anthropogenic pollution, particularly in the form of toxic metals and other elements, and persistent organic pollutants (POPs). Environmental exposure to metals and elements has long been established as of potential toxicological concern in wildlife. These include elements which are essential at low concentrations but toxic if they exceed these (*e*.*g*. copper and iron), as well as non-essential elements that are toxic even at low concentrations (*e*.*g*. lead, cadmium and mercury). In addition to elemental pollutants, an emerging group of organic pollutants that are known to be particularly prevalent in wastewater, are per- and poly-fluoroalkyl substances (PFASs) (Szabo et al., 2023), some of which are also listed or proposed for listing as POPs (UNEP, 2022). Consisting of partially or fully fluorinated carbon chains of varying lengths, these molecules have both hydro- and lipophobic properties (Buck et al., 2011), resulting in their versatile use in *e*.*g*. mining, pesticides, fire-fighting foam, clothing and cosmetics. Some estimates suggest there may be over 14,000 different PFASs marketed globally (US Environmental

Protection Agency, 2022). The strength of the C-F bond in these compounds also contributes to their environmental persistence and biomagnification (Grønnestad et al., 2019; Kelly et al., 2009), which increases with the length of the carbon chain (≥7 C atoms) (Martin, Mabury, Solomon, & Muir, 2003). Moreover, many PFAS precursors can be bio-transformed into particularly persistent compounds in the environment (Wang, Dewitt, Higgins, & Cousins, 2017), which can result in higher concentrations of PFAS in wastewater effluent than inputs would suggest (Gallen, Eaglesham, Drage, Nguyen, & Mueller, 2018). Consequently, PFAS have become widespread in wildlife (AMAP, 2021; Lee, Lee, Lim, & Moon, 2020).

The effects of both elemental pollutants and PFASs on wildlife can be diverse, and include reduced food availability (De Vries et al., 2017), reduced hatching success (Custer et al., 2014), possible endocrine disruption (Ask et al., 2021; Sebastiano et al., 2021), immunomodulation and immunotoxicity (Castaño-Ortiz, Jaspers, & Waugh, 2019; McDonough et al., 2020; Mitra et al., 2022; Vallverdú-Coll, Mateo, Mougeot, & Ortiz-Santaliestra, 2019), and oxidative stress to DNA and proteins in the body (Marasco & Costantini, 2016; Metcalfe & Alonso-Alvarez, 2010; Rainio & Eeva, 2010), leading to tissue damage (Valavanidis, Vlahogianni, Dassenakis, & Scoullos, 2006). One particular PFAS, perfluorooctanesulfonic acid (PFOS), has also been shown to cause immunomodulation in animals, interfering with signalling pathways in the animals’ immune responses (Castaño-Ortiz et al., 2019; Hansen et al., 2020).

Birds are able to excrete pollutants via e.g. faecal matter (Dauwe, Bervoets, Blust, Pinxten, & Eens, 2000), and deposition of pollutants in growing feathers and eggs (Briels et al., 2019; Burger & Gochfeld, 2004; Hargreaves, Whiteside, & Gilchrist, 2010; Løseth et al., 2019). The predominant concern is the tendency of these pollutants to bioaccumulate, that is, when contaminant uptake is higher than its excretion (Berglund, Koivula, & Eeva, 2011; Blomqvist, Frank, & Petersson, 1987). This process can result in accumulation of high concentrations of pollutants in organism tissues and organs, which may threaten wildlife health (Carneiro et al., 2018; Dietz et al., 2019). Hargreaves et al. (2010) examined 17 different elements present in Arctic-breeding shorebirds and found that Hg was elevated to potentially harmful concentrations. Aside from this study, mercury has also been found to occur in harmful concentrations in *Charadrius* plovers (Picone et al., 2019; Su et al., 2020), underscoring the need for more extensive pollutant-burden analyses in shorebirds (Ma, Choi, Thomas, & Gibson, 2022). This is especially necessary in shorebirds as they may be more susceptible to pollution due to the physical stresses from migration (Liess, Foit, Knillmann, Schäfer, & Liess, 2016), and their propensity to forage in soft coastal sediments where pollutants can accrue at much higher concentrations than the surrounding water (Stark, 1998).

Of special concern are effects of pollutants on immune function in avian migrants, given they are known hosts and dispersers of zoonoses such as avian influenza virus (AIV) (Verhagen, Herfst, & Fouchier, 2015; Wille et al., 2019), as well as many other pathogenic microorganisms (Hubálek, 2004). Infection with AIV has been shown to be indicative of high viral diversity in shorebirds (Wille et al., 2018), and may vary with immunomodulatory effects from e.g. PFAS contamination. Whether this affects shorebirds that increasingly utilise artificial wetlands is less well known and would be vital to understand given the increasing reliance of some shorebird populations on artificial wetlands.

To study whether wastewater treatment plants could be a viable alternative habitat to migratory shorebirds, we analysed red blood cell samples from two species of migratory shorebirds commonly found around Australian coasts during the austral summers from 2011 to 2021, at two different sites – one a wastewater treatment plant, another a natural coastal wetland. Our species, curlew sandpipers (*Calidris ferruginea*) and red-necked stints (*Calidris ruficollis*), are both small, long-distance migratory shorebirds that breed in the high Siberian Arctic and are threatened by habitat loss along the flyway (Clemens et al., 2016; Melville, Chen, & Ma, 2016). In addition, ongoing field studies of both species at both sites have also accrued over 40 years of banding data that allow us to conduct survival analyses for insight into population dynamics of these species at our study sites. In both sites and species, we studied concentrations of 15 inorganic elements, 15 PFASs, two oxidative stress (OS) biomarkers, AIV infection and serostatus as well as population survival. Together, these metrics were used to evaluate the adequacy of wastewater treatment plants as suitable alternative habitat for shorebirds and assess the occurrence and effect of pollution and disease in these birds at both an individual and a population level, based on their choice of habitat.

## Methods

### Ethics statement

This research was conducted under approval of Deakin University Animal Ethics Committee (permit numbers A113-2010, B37-2013, B43-2016), Philip Island Nature Park Animal Ethics Committee (SPFL20082). Banding was done under Australian Bird Banding Scheme permit (banding authority numbers 2703, 2915, 8000, 8001)

### Sample Collection

Curlew sandpipers and red-necked stints were captured in Victoria (south-eastern Australia), distributed across 8 major sites along the coast. The two sites where most birds were captured in the course of an annual monitoring program by the Victorian Wader Study Group (VWSG) were chosen as our study sites (Fig. 1 A). The first of these is Melbourne’s Western Treatment Plant (WTP, -37.99 S, 144.62 E), run by Melbourne Water. These wastewater ponds serve to treat the effluent from over half of the population of Melbourne (∼5 million people). They are also renowned for the birdlife that inhabits the area, with considerable efforts made in their management to maintain habitat for wildlife (Steele & Harrow, 2014). Curlew sandpipers and red-necked stints at the WTP carry out much of their foraging at low tide on adjacent flats, which have been enriched by effluent discharges since the 1930s (Steele & Harrow, 2014); at high tide they roost and sometimes forage in constructed, shallow freshwater ponds – mainly former sewage ponds, in which the water supply is now partially or fully treated effluent. For comparison to the WTP, our second sample site was Yallock Creek (−38.21 S, 145.397 E), situated on the northern coast of Western Port Bay to the east of Melbourne. Yallock Creek represents a substantially less-modified coastal wetland, where shorebirds forage on natural tidal flats, and roost above the tideline in settings (e.g. beaches, spits) where little or no supratidal foraging is possible. Distance between these two sites is approximately 80km. Birds were captured by cannon-net as part of long term monitoring program, initiated in 1978 (Minton, 2006). Catches at each site generally occurred one to three times per year, and when caught, birds were banded with an Australian Bat and Bird Banding Scheme band and aged into categories of 1 year (or younger, hereby referred to as ‘immature’), and 2+ year old categories based on plumage characteristics. Blood samples used in all analyses were collected from these birds between 2011 and 2021.

**Figure 1.**
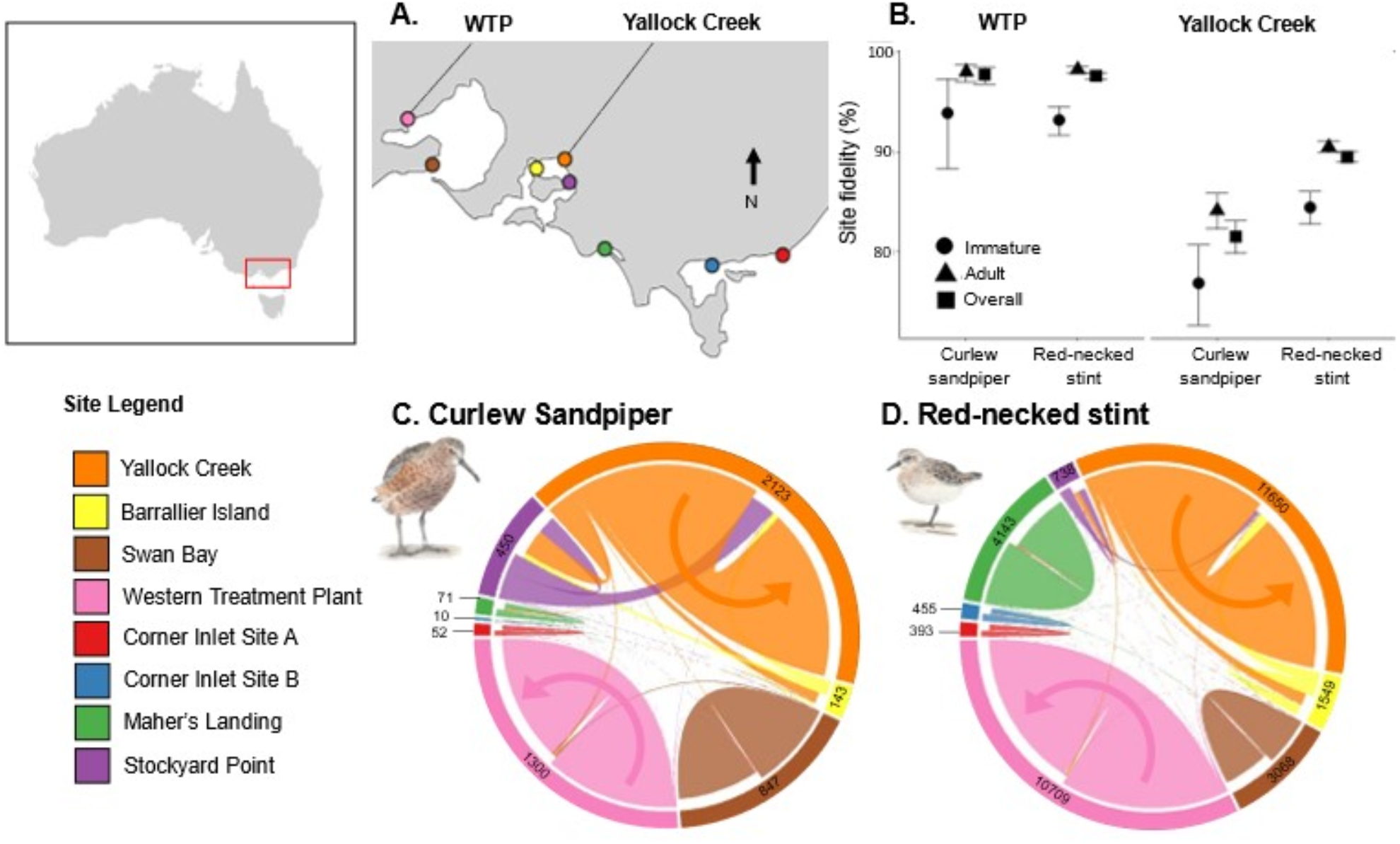
Site fidelity trends in migratory shorebirds captured since 1974 at 8 sites in Victoria, Australia, with the most-sampled sites being WTP and Yallock Creek (A). (B) shows immature, adult and overall site fidelity at both WTP and Yallock Creek. Colours in A correspond to those in C and D. Curlew sandpiper (C) and red-necked stint (D) site fidelities are shown whereby the width of each arc indicates the number of birds first captured at a site (base of arrow, closest to outer ring), and where they were recaptured again (head of arrow, further from outer ring). Bird icons by the author (Tobias Ross).

Blood and swab samples were collected as part of a long term study on avian influenza (Wille et al., 2023). Approximately 200 μl of whole blood was taken from each individual bird from the brachial vein using Microvette capillary tubes (Sarstedt), using new, clean tubes for each individual bird to minimise chance of contamination in the field. These were left to clot and then centrifuged to separate red blood cells (RBC) from serum and stored at <-20 °C. Oropharyngeal and cloacal swabs were taken from each bird, using sterile swabs (Copan). Once collected, swabs from each bird were stored in virus transport media (VTM, brain heart infusion [BHI] broth-based medium [Oxoid] with 0.3 mg/ml penicillin, 5 mg/ml streptomycin, 0.1 mg/ml gentamicin, and 2.5 g/ml, amphotericin B), and stored at -80 °C.

In total, 100 red-necked stint and 70 curlew sandpiper RBC samples were used across all pollutant and OS biomarker analyses. A subset of these was used for each analysis, as due to limitations of sample volume not all samples could be analysed for every pollutant class or OS biomarker.

### Chemical/pollutant analysis

We analysed 86 RBC samples from 2011-2020 for elements, selecting samples to ensure an even representation of years and age groups (immature vs. adults, 43 of each) amongst samples. For these samples, we targeted 15 different elements in our analyses (As, Cd, Co, Cr, Cu, Fe, Hg, Mg, Mn, Ni, P, Pb, Se, Sr, Zn). Sample digestion was performed at the Norwegian University of Science and Technology (NTNU) according to previously published methods with minor modifications (Jantawongsri et al., 2021). Approximately, 0.040 g (±0.028 g) of RBCs were transferred into pre-washed Teflon® tubes (15 mL) and 1.5 mL concentrated ultra-pure grade nitric acid [purified from HNO_3_ (AnalaR NORMAPUR®, VWR) in sub-boiling distillation system, Milestone, SubPur, Sorisole, BG, Italy] was added for a subsequent digestion in a high-pressure microwave system (Milestone UltraCLAVE, EMLS, Leutkirch, Germany) with a maximum temperature of 245 °C at 110 bar for 2.5 h. After digestion, the RBC samples were diluted with ultrapure water (Q-option, Elga Labwater, Veolia Water Systems LTD, UK) to attain a final weight of approximately 35 g (35.61 ± 0.16 g) in 50 mL polypropylene (FalconTM) tubes. Final elemental determination was performed using a triple quadrupole inductively coupled plasma mass spectrometry instrument with ^103^Rh, ^185^Re and ^193^Ir as internal standards (Inorganic Ventures, USA). The analyses were conducted in Norway at SINTEF research institute with Agilent 8900 ICPQQQ (Agilent Technologies, USA) instrument in 2020 and at NTNU (Agilent 8800 ICPQQQ) in 2021.

Three blanks were added in every digestion sample batch and subsequently, all concentrations were corrected for the blank samples. Method detection limits (MDLs) were calculated using three times the standard deviation of the background equivalent concentrations (BEC: blank counts divided by the slope of the calibration curve). To assure quality of analysis, three replicates (one per sample batch) of the quality control material (human whole blood Seronorm Trace Elements Whole BloodTM, Sero, Norway) were prepared. The concentrations found were within 82.2-110.5% of the analytical values, except for As (72.2%).

The concentrations of 15 different PFASs were analysed at NTNU from 150 samples taken between 2011 and 2020, again ensuring an even spread of years and age groups (n= 80 juveniles and 70 adults). The target compounds were broadly classed into three categories, carboxylates (PFPA, PFHxA, PFHpA, PFOA, PFNA, PFDA, PFUnA, PFDoA PFTrA, PFTeA, full names and details of chemicals provided in Table S1), sulfonates (PFBS, PFHxS, PFOS), and total PFASs (including all the aforementioned compounds, as well as PFOSA and NEtFOSA). The RBC samples were extracted using the method of Trimmel et al. (2021) with minor modifications, using hybrid-SPE. The extracts were transferred to amber vials for UPLC-MS/MS analysis. All samples were spiked with ^13^C-isotope labeled PFOS and PFOA IS-mixture. Recoveries ranged from 62% to 157%. A laboratory blank was included in each batch of 20 samples (see Table S2).

### Avian influenza analyses

Samples were screened for avian influenza as per Wille et al. (2021). Briefly, for swab samples collected between 2011 to 2015, RNA was extracted using the MagMax 96 Viral Isolation Kit (Ambio, Thermo Fisher Scientific) using the Kingfisher Flex platform (Thermo Fisher Scientific). RNA was assayed for a short fragment of the matrix gene (Fouchier et al., 2000). First, using the Superscript III Platinum ONE step qPCR Kit (Life Technologies, Thermo Fisher Scientific) with ROX, followed by a subsequent amplification and detection using the SYBR Green mastermix (Life Technologies, Thermo Fisher Scientific). For all samples after 2015, RNA was extracted using the NucleoMag Vet Kit (Scientifix) on the Kingfisher Flex System. Extracted RNA was subsequently assayed for a short fragment of the matrix gene (Spackman et al., 2002) using the SensiFAST Probe Lo-Rox qPCR Kit (Bioline). A cycle threshold (Ct) cut-off of 40 was used. All serum samples were assayed using the Multi Screen Avian Influenza Virus Antibody Test Kit (IDEXX, Hoppendorf, The Netherlands) following manufacturer’s instructions.

### Oxidative Stress Biomarker analyses

Following our pollutant analyses, we utilised oxidative stress biomarkers taken from a subset of our total sample collection, totalling 56 RBC samples. These biomarkers were used as an indicator of the relative health of individual birds at each site, regarding their contamination. To measure the OS of each bird, we chose 8-hydroxy-2-deoxyguanosine (8-OHdG) which is the product of oxidation of DNA, and o,o’-dityrosine (DIY) which indicates peroxidation of proteins (Martinez & Kannan, 2018). The RBC samples were extracted and analysed at NTNU based on the procedure reported by Zhang et al. (2022). All samples were spiked with 8-OHDG-^15^N_5_ and DIY-^13^C_12_. Recoveries of 8OHDG and DIY were 101% and 96%, respectively.

### Statistical analyses

#### Site fidelity

We used long-term banding data from 1978 to 2019 to highlight movement between the sampling sites (primarily WTP and Yallock Creek). Exchange between sites was evaluated using each individual bird’s recapture event, taking into account where they were first captured. We made three site fidelity analyses, focusing on i) immature site fidelity, ii) adult site fidelity, and iii) overall population site fidelity. The significance of any difference between immature and adult site fidelity was determined using an exact binomial test in R. Data for overall population site fidelity were depicted using the package circulize (Version 0.4.15) implemented in R version 4.2.0 (Gu, Gu, Eils, Schlesner, & Brors, 2014).

#### Elements, PFAS, AIV and oxidative stress biomarker analyses

All statistical analyses were conducted using R. All 15 PFASs were summed into their respective categories. Where a compound/group of compounds (out of 15 elemental pollutants, Total PFASs, PFCAs, and PFSAs) was not detected in 50% or more of the samples, this was excluded from statistical analyses. For each pollutant as well as each biomarker we investigated the effects of species, age group (immature vs adult) and sample site using linear mixed effect modelling from R package lme4 (Bates, Mächler, Bolker, & Walker, 2015) with year as a random factor. For all elements, summed PFAS categories, and oxidative stress biomarkers where the sample concentrations were below the level of detection (LOD), concentrations were set at 0.015 ng/g (half of the lowest LOD in the study) prior to statistical analysis. Additionally, all PFAS concentrations were log transformed to remove skewness and guarantee homogeneity of variances in the data. To analyse viral and sero-prevalence of AIV we used generalised linear mixed-effect models with family set to binomial, with dependent variables of region and age group.

#### Survival analyses

The banding data from 1978 to 2019 data was used to estimate survival of each species at each sampling site, using Bayesian population analyses adapted from Kéry and Schaub (2011). Each of these four models were conducted using WinBUGS14, and R within RStudio. In each model we assumed a constant, age-dependent (*i*.*e*. 1 and 2+) annual survival (Φ) across the 40-year study period, i.e. each model led to the estimation of an average immature and an average adult survival rate. We furthermore assumed a fixed recapture rate (*p*) for both age classes but variable across years *(i*.*e*. for each year of the study a separate recapture rate was estimated to accommodate for annual variations in capture effort). Models were run using four Markov chain Monte Carlo (MCMC) simulations of 30,000 iterations each, discarding the first 15000 and using a thinning factor of 20 to result in 3000 iterations being saved, which were used to describe the posterior distribution of each parameter (in this case, mean adult and immature survival and annual recapture rate and the 95% confidence interval of each). To evaluate the accuracy of Φ and *p* estimates, we used 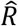 and ensured that all models had 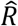 values as close to 1 as possible (for all cases, 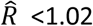 <1.02).

## Results

### Site fidelity

In order to confirm that the sites we used in our study were independent with limited bird movement, we performed a site fidelity analysis. Over the 40-year VWSG banding period, a total of 4995 curlew sandpipers and 32,705 red-necked stints recaptures were recorded across all catching sites, unequivocally showing that site faithfulness to WTP and Yallock Creek was very high (Figure 1A-D). Binomial tests indicated that curlew sandpipers demonstrated 97.8% (95% CI: 96.8-98.5) and 81.5% (95% CI: 79.8-83.1) fidelity to WTP and Yallock Creek and red-necked stints showed 97.6% (97.3-97.9) and 89.5% (88.9-90.0) fidelity, to both sites respectively. When broken down into immature and adult constituents, immature curlew sandpipers showed 93.9% (88.3-97.3) and 76.8% (72.6-80.1) site fidelity to WTP and Yallock Creek, and immature red-necked stints showed 93.2% (91.7-94.5) and 84.4% (82.8-86.0) fidelity to both sites respectively. This is compared with adult birds, where curlew sandpipers showed 98.1% (97.0-98.8) and 84.1% (82.3-85.8), while red-necked stints showed 98.2% (98.0-98.5) and 90.5% (89.9-91.1) to each site respectively. The lower site faithfulness to Yallock Creek compared to WTP was likely due to a higher availability of alternative sites in Western Port Bay and if these sites were included we would likely see higher site fidelity within Western Port Bay than just at Yallock Creek. However, as we only have analysed samples from Yallock Creek (the largest catch site on Western Port Bay), we have limited our site fidelity analyses to this specific site only. The overall curlew sandpiper exchange rate from WTP to Yallock Creek was 0.5% and the reverse exchange rate was 0.9%. For red-necked stints, overall rates were 0.7% and 0.8%. Despite no account of annual variation in catching effort, this quantity of data over 40 years overwhelmingly shows the vast majority of birds spent most, if not all of their non-breeding seasons in either WTP or Yallock Creek.

### Element and PFAS concentrations

Four compounds, chromium (Cr), cobalt (Co), nickel (Ni) and cadmium (Cd) were not detected in more than 50% of samples and so were not included in statistical analyses. Of the 11 remaining elements, 8 showed no significant difference between species, regions or age groups (Table 1, Figure 2, Tables S3 and 4). Three elements exhibited significant difference with at least one explanatory variable (Table 1). Specifically, Iron (Fe) was found to be significantly lower in red-necked stints than in curlew sandpipers (Figure 2B), as was selenium (Se). Se was also lower at the Western Treatment Plant than Yallock Creek (Figure 2C; WTP median: 5000 ng/g, range: 1950-34400 ng/g; Yallock Creek median: 19200 ng/g, range: 4130-65200 ng/g), and lower in immature birds than adults. By contrast, mercury was found to be higher in the WTP than Yallock Creek (Figure 2D, WTP median: 224 ng/g, range: 19-873 ng/g; Yallock Creek median: 160 ng/g, range: 22-998 ng/g).

**Table 1.**
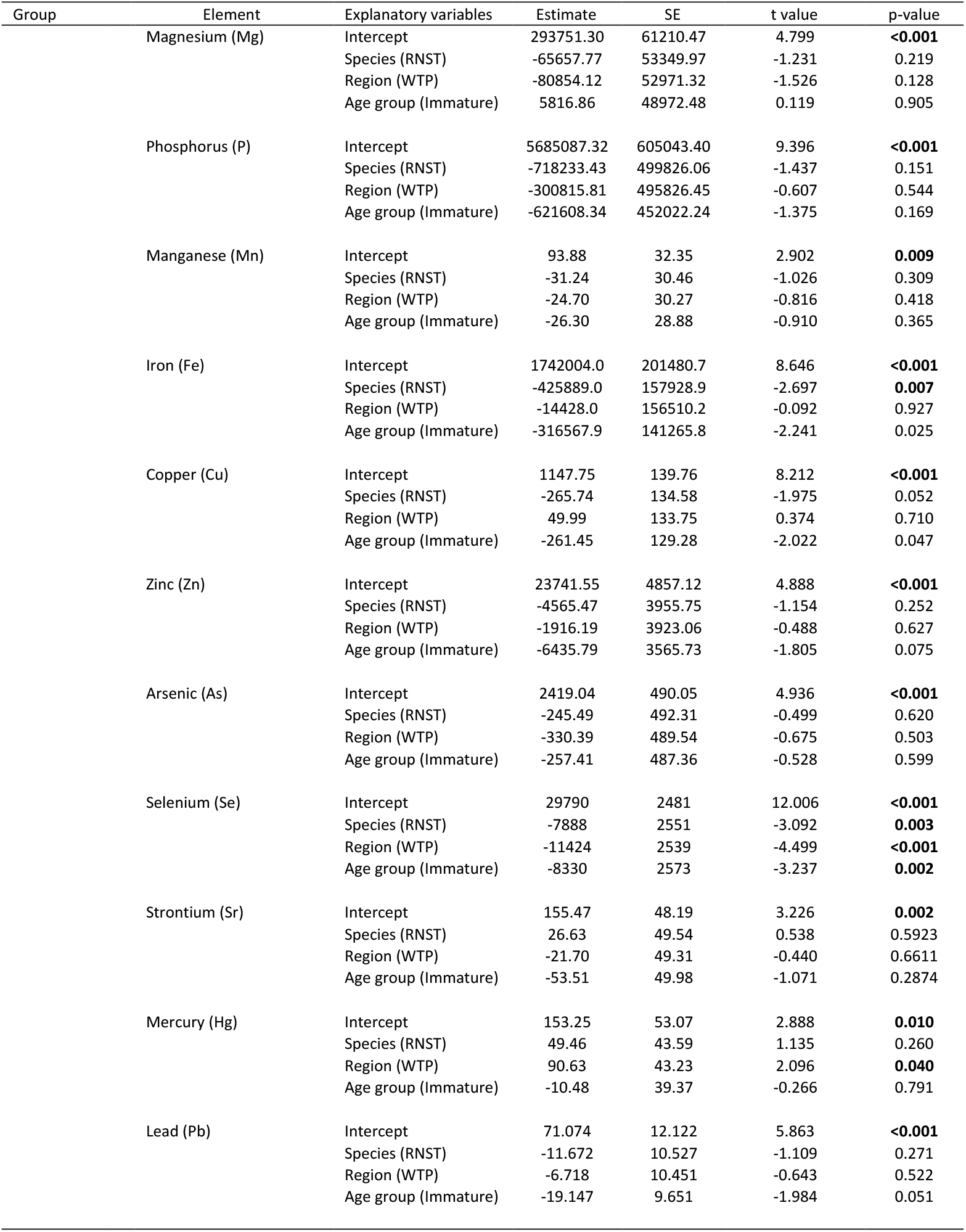
Linear mixed effect model summaries for of each elemental pollutant with detection rate above 50% of samples. Mixed effects models include species, region and age class (adult vs immature) as fixed effects, and sample year as a random effect.

**Figure 2.**
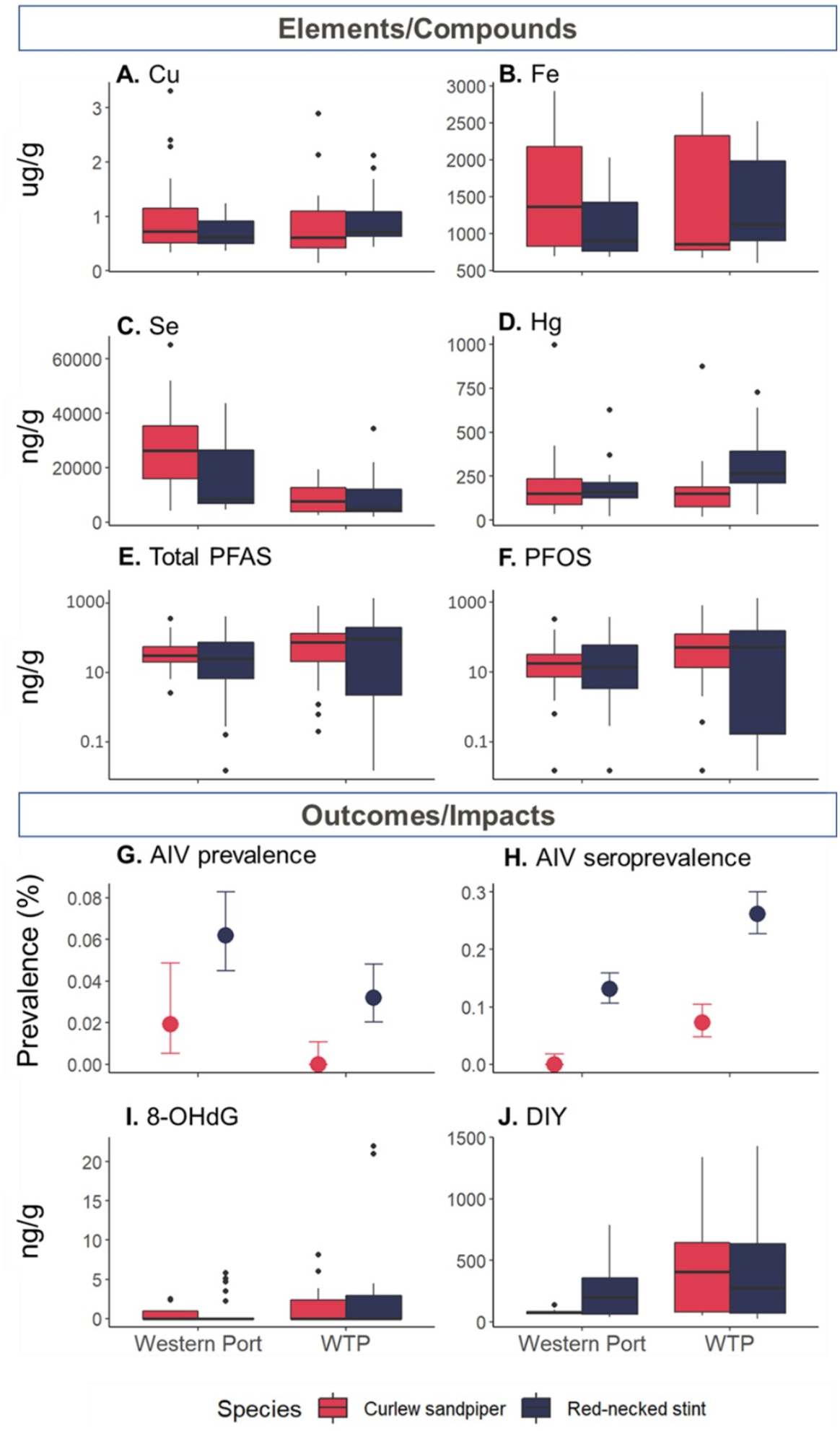
Pollutant, oxidative stress biomarker and avian influenza (AIV) data for each species at each sample site, Yallock Creek which is situated in Western Port Bay, and Melbourne’s Western Treatment Plant (WTP). Contaminants displayed in each plot are as follows (only elements of note are shown to illustrate trends or lack thereof): A: copper (Cu), B: iron (Fe), C: selenium (Se), D: mercury (Hg), E: Total PFAS, F: PFOS, G: avian influenza virus prevalence, H) AIV seroprevalence, I) 8-0HdG biomarker, J) DIY biomarker. Significant species variations shown in B and C, significant region differences in C, E, and J.. Variation between adult and immature age groups not shown. All other contaminants not displayed here are provided in Table S4.

No significant differences were discerned in the mean concentrations of ∑PFCAs or ∑PFSAs between species, region or age group (Table 2). However, the total PFASs concentrations we observed were found to have a statistically significant difference between region and age group, with higher concentrations observed at the WTP (median: 71 ng/g; range: <0.01-1360 ng/g) than Yallock Creek (median 26 ng/g; range: <0.01-414 ng/g), and lower concentrations in immatures than in adults (Figure 2E). This pattern was driven by the most prevalent individual PFAS in our study, PFOS, which also exhibited the same significant patterns between region and age group (Figure 2F, WTP median: 52 ng/g, range: <0.01-1280 ng/g; Yallock Creek median: 14 ng/g, range: <0.01-379 ng/g), and likely was the driver of the significant difference in total PFASs.

**Table 2.** Model summaries for linear mixed effects models of PFASs groups (total PFAS, total carboxylates/PFCAs, total sulfonates/PFSAs, and PFOs by itself), and oxidative stress biomarkers, and general linear models for avian influenza prevalence data. In linear mixed effects models, PFASs were log-transformed to guarantee homogeneity in variances, and models for both PFASs and biomarkers included species, region and age group (adult vs immature) as fixed effects, and sample year as a random effect. Glms for avian influenza had binomial distributions, and were run only for red-necked stints, including only region and age-group as a explanatory variables.

| Group | Model | Dependent variable | Explanatory variables | Estimate | SE | t value/z-value | p-value |
| --- | --- | --- | --- | --- | --- | --- | --- |
| All grouped PFAS (log-transformed) | Linear mixed effects models (lmer) | ΣPFAS (log transformed) | Intercept | 1.253 | 0.511 | 2.452 | **0.029** |
|  |  |  | Species (RNST) | -0.304 | 0.270 | -1.127 | 0.262 |
|  |  |  | Region (WTP) | 0.534 | 0.258 | 2.068 | **0.040** |
|  |  |  | Age group (Immature) | -0.653 | 0.253 | -2.583 | **0.011** |
|  |  | ΣPFCAs (log-transformed) | Intercept | -0.193 | 0.561 | -0.343 | 0.735 |
|  |  |  | Species (RNST) | -0.279 | 0.440 | -0.635 | 0.527 |
|  |  |  | Region (WTP) | 0.558 | 0.424 | 1.318 | 0.189 |
|  |  |  | Age group (Immature) | -0.724 | 0.423 | -1.718 | 0.088 |
|  |  | ΣPFSAs (log-transformed) | Intercept | 0.678 | 0.817 | 0.830 | 0.423 |
|  |  |  | Species (RNST) | -0.336 | 0.388 | -0.865 | 0.388 |
|  |  |  | Region (WTP) | 0.350 | 0.372 | 0.940 | 0.349 |
|  |  |  | Age group (Immature) | -0.653 | 0.363 | -1.799 | 0.074 |
|  |  | PFOS (log-transformed) | Intercept | 1.018 | 0.450 | 2.261 | **-0.040** |
|  |  |  | Species (RNST) | -0.351 | 0.249 | -1.410 | 0.161 |
|  |  |  | Region (WTP) | 0.562 | 0.239 | 2.353 | **0.020** |
|  |  |  | Age group (Immature) | -0.528 | 0.234 | -2.257 | **0.026** |
| Avian influenza virus (AIV) prevalence (glm models) | Generalised linear models (glm) | AIV prevalence | Intercept | -2.834 | 0.180 | -15.783 | **<0.001** |
|  |  |  | Site (WTP) | -0.687 | 0.269 | -2.551 | **0.012** |
|  |  |  | Age group (Immature) | 0.459 | 0.283 | 1.624 | 0.104 |
|  |  | AIV seroprevalence | Intercept | -1.673 | 0.119 | -14.091 | **<0.001** |
|  |  |  | Site (WTP) | 0.848 | 0.152 | 5.598 | **<0.001** |
|  |  |  | Age group (Immature) | -1.564 | 0.279 | -5.621 | **<0.001** |
| Oxidative stress biomarkers | Linear mixed effects models (lmer) | 8-OHdG | Intercept | -0.611 | 1.512 | -0.404 | 0.690 |
|  |  |  | Species (RNST) | 1.573 | 1.082 | 1.454 | 0.150 |
|  |  |  | Region (WTP) | 1.607 | 0.845 | 1.901 | 0.061 |
|  |  |  | Age group (Immature) | 1.429 | 0.967 | 1.478 | 0.144 |
|  |  | DIY | Intercept | 261.76 | 133.28 | 1.964 | 0.064 |
|  |  |  | Species (RNST) | -48.41 | 90.30 | -0.536 | 0.593 |
|  |  |  | Region (WTP) | 224.29 | 69.80 | 3.213 | **0.002** |
|  |  |  | Age group (Immature) | -60.99 | 80.04 | -0.762 | 0.449 |

### Pollutants associations to disease and oxidative stress

As metals, other elements and PFAS may modulate the immune systems of birds, we used avian influenza infection as a disease marker in this study. A total of 215 curlew sandpiper and 855 red-necked stint samples from Yallock Creek, and 430 and 775 from the same respective species at the WTP were used to determine AIV prevalence and seroprevalence. Red-necked stints from Yallock Creek were found to have higher rates of active infection than stints from the WTP (Figure 2G), however, stints from the WTP had significantly higher rates of seropositivity than their Yallock Creek counterparts (Figure 2H). As infection status and seroprevalence are modulated by bird age, we tested whether this pattern was driven by age. It does not appear to be, as no significant difference in active infection status was detected between age groups (adults vs immatures), however immature birds had significantly lower rates of seropositivity than adults (Table 2).

We further investigated markers of oxidative stress in relation to metals, elements and PFAS. No significant difference was found between concentrations in either species or age group for either of our biomarkers (Table 2), and no difference between regions for 8-OHdG (Figure 2I). However, we did find a significant difference between regions for DIY, with slightly higher average concentrations observed in the individuals at the WTP than Yallock Creek (Figure 2J).

### Apparent survival

For both curlew sandpiper and red-necked stint, local survival analyses yielded no significant differences (i.e. 95% CI’s were overlapping) in adult survival between the two sites (see Table 3 for summary). Adult curlew sandpiper apparent survival was 70.5% (95% CI: 68.8-72.1%) and 71.7% (70.3-72.9%) at WTP and Yallock Creek respectively, while red-necked stint survival was 69.3% (68.7-69.8%) and 66.7% (66.0-67.4%) at WTP and Yallock Creek respectively. However, one particularly interesting finding is that we did find significant lower apparent survival of immature birds at WTP than at Yallock Creek in both curlew sandpiper (WTP 61.2% [50.1-73.0%]; Yallock Creek 82.1% [73.4-91.4%]) and red-necked stint (WTP 63.6% [59.9-67.4%]; Yallock Creek 75.3% [72.1-78.5%]). In both species the average immature survival was higher than adult survival in Yallock Creek, but lower in WTP (Figure 3), which was significantly different for red-necked stints. However, in curlew sandpipers this difference was non-significant due to overlapping confidence intervals.

**Table 3.** Summary of site fidelity results for each species, showing fidelities specific to each site for each age group. 95% confidence intervals are shown in brackets.

| Species | Site | Adults | Immatures |
| --- | --- | --- | --- |
| Curlew Sandpiper | WTP | 70.5% (68.8-72.1%) | 61.2% [50.1-73.0%] |
|  | Yallock Creek | 71.7% (70.3-72.9%) | 82.1% [73.4-91.4%] |
| Red-necked Stint | WTP | 69.3% (68.7-69.8%) | 63.6% [59.9-67.4%]; |
|  | Yallock Creek | 66.7% (66.0-67.4%) | 75.3% [72.1-78.5%] |

**Figure 3.**
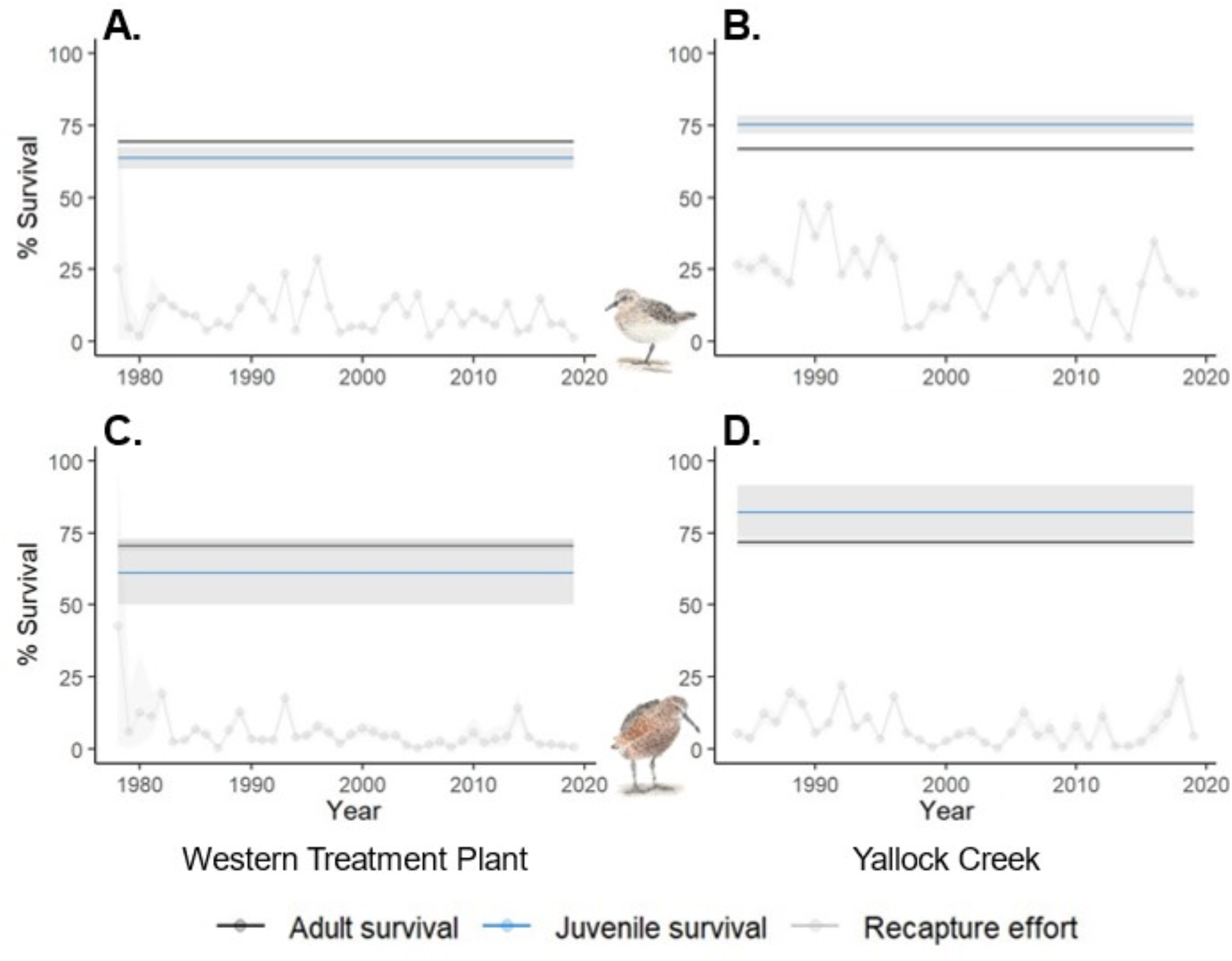
Average survival of adult and immature red-necked stints (A and B)) and curlew sandpipers (C and D) at WTP and Yallock Creek on Western Port Bay, from 1978-2019 for WTP samples and 1984-2019 for YC samples. Grey illustrates confidence intervals surrounding the average survival of1ach age group. Bird icons by the author (Tobias Ross).

## Discussion

Although a waste-water treatment plant may have the connotation of being a potentially suboptimal habitat compared to a natural habitat, our collective results, encompassing the analysis of as many as 30 pollutants, two oxidative stress biomarkers, a key disease marker and a survival analysis for two shorebird species, do not suggest the investigated WTP to be an inferior habitat for shorebirds compared to Yallock Creek.

### Site fidelity

Our analyses showed high site fidelity of these shorebirds, indicating that our local survival data on adult is unlikely to be heavily influenced by the emigration of birds from the WTP (where the VWSG has caught birds) to other sites (where birds are not caught, e.g. inland). Our local adult survival estimates for both species were broadly comparable to those presented by Méndez, Alves, Gill, and Gunnarsson (2018), differing only by 2-4% in total, and our indication of minimal exchange between sites once again highlighted the extreme site fidelity shown by many migratory shorebirds (Christie & Jessop, 2009; Coleman & Milton, 2012; Warnock & Takekawa, 2008). Such high site fidelity allowed us confidence that differences seen in exposure between sampling sites were indeed reflective of the contamination in these different environments.

### Inorganic element and PFAS concentrations

Hg and PFOS concentrations were the only two pollutants of our 30 targeted elements/compounds that showed any indication that the WTP is a more polluted environment than the relatively unmodified coastal wetland of Yallock Creek. No other pollutants showed no significant differences between sites, except for Se which was found to be higher at Yallock Creek than WTP. In this case, Se at Yallock Creek may be attributable to geogenic sources in the region rather than anthropogenic sources (Gebreeyessus & Zewge, 2019). This may have different reasons such as different elemental composition of the soil, minerals in rock and the bioavailability of different elemental forms (speciation) in different locations such as presented for the case with Se (Winkel et al., 2012). In addition, agriculture could present a source of different elements to the environment, such as through the use of fertilisers and pesticides (Zhang & Wang, 2020). Where there are significant differences between age-group concentrations (seen in Se and PFOS/total PFASs), this can likely be attributed to the tendency of both to bioaccumulate during the birds’ life spans; immature birds have not had the continued exposure to garner high concentrations that an adult may have (Lee et al., 2020; Yue, Huang, Wang, & Qiao, 2021). It is possible that the minimal differences in elements and PFASs concentrations that we observed between sites were also due to the proximity of Yallock Creek to the Melbourne region, hence it may not be as clean as sites more remote from human settlements.

Moreover, our contaminant concentrations in both sites also appeared generally low, with only two main exceptions (As and Se) appearing to exceed possible adverse effects levels in blood (Burger et al., 2019; Tsipoura et al., 2017). This is also supported by our biomarker 8-OHdG for oxidative damage to DNA where we saw no difference between sampling sites, and nor did we see any difference in the survival of adults. We did, however, see differences in apparent local survival of first year birds between sites (addressed below), as well as some difference in AIV infection load in the population where birds at the WTP showed less prevalence of active AIV infections than birds at Yallock Creek. However, birds at the WTP had higher seroprevalence, which suggested that these individuals had been previously exposed to AIV more frequently compared to Yallock Creek birds. This observation paired with the higher Hg and PFOS concentrations we saw at the WTP, could contribute to the higher levels of oxidative damage to proteins indicated by our DIY indicator at the WTP. Despite this last finding, the overall low concentrations and minimal differences between regions that we observed tentatively suggest that appropriately managed wastewater treatment wetlands may be suitable alternative habitat for shorebirds and may potentially compensate for habitat loss.

Our suggestion of the use of wastewater wetlands as novel habitat is not without caveats. The higher concentrations of PFOS (and therefore total PFASs) and mercury at the WTP certainly warrant some concern; indeed sewage in wastewater plants is an established environmental source of PFASs (Arvaniti & Stasinakis, 2015) and elements (Farkas et al., 2020), and so may yet threaten wildlife. The absolute concentrations of pollutants seen in both red-necked stints and curlew sandpipers might still be of concern for conservation outlooks, especially given that pollutants such as these have been noted to be persistent in animal tissues (Stahl, Mattern, & Brunn, 2011). While our samples had lower concentrations of most elemental pollutants than observed in other studies of elements in whole blood of shorebirds (Burger et al., 2019; Hargreaves et al., 2010; Tsipoura et al., 2017) in which such levels were noted to be lower than toxicological thresholds, our As and Se concentrations were found to greatly exceed those seen in Burger et al. (2019), and Hargreaves et al. (2010). In the case of the latter of these two studies, our Hg concentrations were lower by comparison, even at the WTP, which could be because our birds were caught when actively moulting, or having just finished moult and thereby excreted Hg into their feathers (Furness, Muirhead, & Woodburn, 1986). The As concentrations at both sites and the Se concentrations at Western Port Bay appeared higher than toxicological thresholds for sublethal effects (Burger et al., 2019; Tsipoura et al., 2017), which is cause for concern for the health of these birds, despite Se being an essential element. By comparison, the Se concentrations seen in the WTP birds likely pose little threat to their health.

Regarding PFASs, our use of RBC differed from the more commonplace use of serum in other PFAS studies (Lopez-Antia et al., 2019; Løseth et al., 2019; Newsted, Jones, Coady, & Giesy, 2005). Gebbink and Letcher (2012) found relationships between plasma and RBC concentrations of PFOS to be approximately 4:1. When this relationship is applied to our collective PFAS data, it suggests that our observed PFASs contamination for all but PFOS in some individuals were predominantly lower than thresholds discussed in Newsted et al. (2005). These concentrations may still pose a threat at sublethal levels, particularly compared to the lowest liver concentrations with observable effects reported in bobwhite quail (*Colinus virginianus*) by Dennis et al. (2021). Comparisons with these liver concentrations are only tentative due to different matrices, but blood concentrations of PFOS tend to be lower than those seen in liver (Gebbink & Letcher, 2012). However, in a small number of samples from both sites, PFOS concentrations were particularly elevated. Occurrence of such elevated concentrations with potential harmful health effects reinforces the importance of ongoing monitoring of pollutants for shorebird conservation, especially at artificial wetlands such as the WTP.

### Impacts on immune response

Our AIV infection data provides a valuable insight into the immunocompetence of our shorebirds. Many pollutants can influence the way birds’ immune systems can respond to pathogens, either by being directly immunotoxic (Mitra et al., 2022) or by modulating aspects of immune response such as altering antibody function and inflammatory responses (Dewitt, Peden-Adams, Keller, & Germolec, 2012). There is some evidence to suggest that migration may also reduce immune function (Eikenaar & Hegemann, 2016), which would be problematic for shorebirds. Along their migration in the EAAF, they traverse an avian influenza hotspot (Wille et al., 2019), infection by which can be used as a marker for other diseases (Wille et al., 2018). Birds with reduced immune function are inevitably more susceptible to disease, and a combination of migration and pollution stress on immune response could directly increase the prevalence of disease in the population, thereby decreasing their population survival.

In our study, we found significant differences in both AIV prevalence and seroprevalence, however, due to no consistent differential pollution between either sample site we cannot attribute these differences to exposure to contaminants. If the WTP was found to be a consistently more polluted site, we would have expected AIV infection data to show patterns similar to those seen in AIV seroprevalence, where individuals at the WTP have higher levels than those at Yallock Creek. This was not the case. However, the reason that red-necked stints at the WTP have lower prevalence of AIV infections than birds inhabiting Yallock Creek remains unknown. By comparison, the higher seroprevalence of AIV in adult birds than immature birds was as expected. Juvenile birds are naíve to the disease by comparison to their older conspecifics, as they have had less time to be exposed and then recover, resulting in lower prevalence of antibodies in their blood serum (Wille et al., 2023).

### Apparent survival

Pollution can present an insurmountable threat to survival of wildlife. Therefore, it was important that we analysed this in our assessment of habitat quality, in which we found no difference in adult survival between locations. There are widespread declines in shorebirds that have been documented along the flyway (Clemens et al., 2016; Studds et al., 2017), but we don’t appear to find any differences between either our species or our sites. This suggests that any effects on adult survival are not site-specific in Australia, and observed population declines are related to other factors in the flyway rather than any differential exposure to pollution due to habitat choice within Australia.

Where we did see differences in pollution concentrations between sampling sites, such as in Hg, PFOS and Se, high site fidelity allowed us confidence that these were indeed reflective of site-related differences. However, our analyses only highlight local survival, and due to difficulties in measuring true survival (e.g. limited by the inability to catch birds in every region of their distribution), true survival is likely higher than our local survival estimates. Nevertheless, the different apparent local survival of immature birds between both sites raised two interesting points. Firstly, immature survival was higher than that of adults at Yallock Creek in both red-necked stints and curlew sandpiper. Immature survival within Australia does not reflect the juvenile birds that perish along their migration; only the strongest juveniles arrive in Australia and it is their survival that is being estimated, causing the appearance of a higher rate of immature survival than adult survival here. Not only this, the immature birds also do not migrate the following year after they first arrive in Australia (Menkhorst et al., 2019), which takes out two potentially arduous long-distance migrations to and from the breeding grounds, compared to their adult con-specifics.

Secondly, immature survival was reduced in the birds from the WTP. Our survival analyses examine local survival rather than true survival and were unable to account for undetected movement beyond our sample sites. This is particularly pertinent to immature survival, as first year juveniles do not migrate the following year from their initial arrival, and may exhibit increased exploratory behaviours and travel widely in this first year of life in Australia (D. Rogers, pers. comm.), which has been posited to be fuelled by an increase in mass over the austral winter despite not migrating (Rogers, Rogers, & Minton, 1996). Such exploration is also seen in other species of shorebirds such as red knots (*Calidris canutus*) and bar-tailed godwits (*Limosa lapponica*) (Battley, Schuckard, & Melville, 2011; Battley et al., 2020). Our sampling efforts on Port Phillip Bay are concentrated at the WTP, but immature birds wander and inhabit other areas nearby that we do not sample thereby ‘removing’ them from our sample population, which is indicated by our slightly lower immature site fidelity (Figure 1B). At Western Port Bay, there are several catching sites within close proximity so this movement can be detected (e.g. purple in Figure 1A, C-D) but with no other catching sites within the WTP, such movement would be lost despite our observed high site fidelity in recaptured immatures. As a result, immatures leaving the WTP for other locations where they are not caught again could account for the apparent lower immature survival seen at this site.

## Considerations and conclusion

Threats other than the pollutants we have analysed may also require further investigation. For example, current-use pesticides from agricultural runoff in sites like Yallock Creek may harm the region’s wildlife (Chen et al., 2020) and contribute to a different pollutant profile when compared to a wastewater plant. On the other hand, emerging pollutants such as pharmaceutical contamination (Patel et al., 2019), and antimicrobial resistance in bacteria in wastewater (Marcelino et al., 2019) may impact birds inhabiting the WTP in ways we have not investigated in this study. Indeed, pharmaceutical contamination and antimicrobial resistance could also contribute to the lower apparent immature survival we have seen. Immature foraging proficiency is often lower than adult birds resulting in differences in strategy and prey choice (Groves, 1978; Puttick, 1979), which might result in higher susceptibility to pollutants such as pharmaceuticals, as well as antimicrobial resistant pathogens in the environment. Additionally, legacy POPs may also pose a threat to shorebirds (Ma et al., 2022), and these have not been included in the current study. More studies are required in these fields about the health of shorebirds to complete our understanding of the effects of wastewater on contamination in shorebirds. Ongoing monitoring is also vital considering elevated PFOS concentrations that we have seen in a small minority of our studied birds. Nevertheless, when taken together, our findings regarding pollutant concentrations, health status, and survival in shorebird populations at both sites indicate negligible differences, and tentatively suggest that controlled wastewater treatment plants could provide a valuable alternative habitat to migratory shorebirds along their flyway.

## Acknowledgements

This project was funded by the Australian Research Council (ARC) Discovery Project Grant DP190101861 (to Marcel Klaassen) and additional funding by the Norwegian Research Council (NRC) project COAST IMPACT 302205/SHJ (to Veerle L. B. Jaspers). The WHO Collaborating Centre for Reference and Research on Influenza is funded by the Australian Department of Health. Michelle Wille is funded by an Australia Research Council Discovery Early Career Research Award (DE200100977). We would also like to thank Melbourne Water, in particular Suelin Haynes, Judy Blackbeard, Philip Wilkie, Nick Crosbie and Will Steele for their role in facilitating all our work in catching birds at the WTP, and for reviewing this manuscript. Thanks also go to Danny Rogers for providing revisions on the manuscript on behalf of the VWSG.

## References

AMAP. (2021). AMAP assessment 2020: POPs and chemicals of emerging arctic concern: influence of climate change (pp. viii+134pp). Tromsø, Norway.: Arctic Monitoring and Assessment Programme (AMAP).

Arvaniti, O. S., & Stasinakis, A. S. (2015). Review on the occurrence, fate and removal of perfluorinated compounds during wastewater treatment. Science of the Total Environment, 524-525, 81–92. doi: https://doi.org/10.1016/j.scitotenv.2015.04.023

Ask, A. V., Jenssen, B. M., Tartu, S., Angelier, F., Chastel, O., & Gabrielsen, G. W. (2021). Per- and polyfluoroalkyl substances are positively associated with thyroid hormones in an arctic seabird. Environmental Toxicology and Chemistry, 40(3), 820–831. doi: 10.1002/etc.4978

Bates, D., Mächler, M., Bolker, B., & Walker, S. (2015). Fitting linear mixed-effects models using lme4. Journal of Statistical Software, 67(1), 1–48. doi: 10.18637/jss.v067.i01

Battley, P., Schuckard, R., & Melville, D. (2011). Movements of bar-tailed godwits and red knots within New Zealand Science for Conservation (Vol. 315).

Battley, P. F., Conklin, J. R., Parody-Merino, Á. M., Langlands, P. A., Southey, I., Burns, T., et al. (2020). Interacting roles of breeding geography and early-life settlement in godwit migration timing. Frontiers in Ecology and Evolution, 8(52). doi: 10.3389/fevo.2020.00052

Berglund, Å. M. M., Koivula, M. J., & Eeva, T. (2011). Species- and age-related variation in metal exposure and accumulation of two passerine bird species. Environmental Pollution, 159(10), 2368–2374. doi: https://doi.org/10.1016/j.envpol.2011.07.001

Blomqvist, S., Frank, A., & Petersson, L. R. (1987). Metals in liver and kidney tissues of autumn-migrating dunlin Calidris alpina and curlew sandpiper Calidris ferruginea staging at the Baltic Sea. Marine Ecology Progress Series, 35(1/2), 1–13.

Briels, N., Torgersen, L. N., Castaño-Ortiz, J. M., Løseth, M. E., Herzke, D., Nygård, T., et al. (2019). Integrated exposure assessment of northern goshawk (Accipiter gentilis) nestlings to legacy and emerging organic pollutants using non-destructive samples. Environmental Research, 178, 108678. doi: 10.1016/j.envres.2019.108678

Buck, R. C., Franklin, J., Berger, U., Conder, J. M., Cousins, I. T., De Voogt, P., et al. (2011). Perfluoroalkyl and polyfluoroalkyl substances in the environment: Terminology, classification, and origins. Integrated Environmental Assessment and Management, 7(4), 513–541. doi: 10.1002/ieam.258

Burger, J., & Gochfeld, M. (2004). Marine birds as sentinels of environmental pollution. EcoHealth, 1(3), 263–274. doi: 10.1007/s10393-004-0096-4

Burger, J., Mizrahi, D., Jeitner, C., Tsipoura, N., Mobley, J., & Gochfeld, M. (2019). Metal and metalloid levels in blood of semipalmated sandpipers (Calidris pusilla) from Brazil, Suriname, and Delaware Bay: Sentinels of exposure to themselves, their prey, and predators that eat them. Environmental Research, 173, 77–86. doi: https://doi.org/10.1016/j.envres.2019.02.048

Carneiro, M., Oliveira, P., Brandão, R., Soeiro, V., Pires, M. J., Lavin, S., et al. (2018). Assessment of the exposure to heavy metals and arsenic in captive and free-living black kites (Milvus migrans) nesting in Portugal. Ecotoxicology and Environmental Safety, 160, 191–196. doi: https://doi.org/10.1016/j.ecoenv.2018.05.040

Castaño-Ortiz, J. M., Jaspers, V. L. B., & Waugh, C. A. (2019). PFOS mediates immunomodulation in an avian cell line that can be mitigated via a virus infection. BMC Veterinary Research, 15(214), 1–9. doi: 10.1186/s12917-019-1953-2

Chen, C., Zou, W., Cui, G., Tian, J., Wang, Y., & Ma, L. (2020). Ecological risk assessment of current-use pesticides in an aquatic system of Shanghai, China. Chemosphere, 257, 1–10. doi: https://doi.org/10.1016/j.chemosphere.2020.127222

Christie, M., & Jessop, R. (2009). Site faithfulness of ruddy turnstone Arenaria interpres in the south east of South Australia (pp. 1–24): Wildlife Conservation Fund Research Grants Programme.

Clemens, R., Rogers, D. I., Hansen, B. D., Gosbell, K., Minton, C. D. T., Straw, P., et al. (2016). Continental-scale decreases in shorebird populations in Australia. Emu, 116(2), 119–135. doi: 10.1071/MU15056

Coleman, J., & Milton, D. (2012). Feeding and roost site fidelity of two migratory shorebirds in Moreton Bay, South-eastern Queensland, Australia. Sunbird, 42, 41–51.

Conklin, J., Verkuil, Y., & Smith, B. (2014). Prioritizing migratory shorebirds for conservation action on the East Asian-Australasian flyway: WWF Hong Kong.

Custer, C. M., Custer, T. W., Dummer, P. M., Etterson, M. A., Thogmartin, W. E., Wu, Q., et al. (2014). Exposure and effects of perfluoroalkyl substances in tree swallows nesting in Minnesota and Wisconsin, USA. Archives of Environmental Contamination and Toxicology, 66(1), 120–138. doi: 10.1007/s00244-013-9934-0

Dauwe, T., Bervoets, L., Blust, R., Pinxten, R., & Eens, M. (2000). Can excrement and feathers of nestling songbirds be used as biomonitors for heavy metal pollution? Archives of Environmental Contamination and Toxicology, 39, 541–546. doi: 10.1007/s002440010138

De Vries, P., Slijkerman, D. M. E., Kwadijk, C. J. A. F., Kotterman, M. J. J., Posthuma, L., De Zwart, D., et al. (2017). The toxic exposure of flamingos to per - and polyfluoroalkyl substances (PFAS) from firefighting foam applications in Bonaire. Marine Pollution Bulletin, 124(1), 102–111. doi: 10.1016/j.marpolbul.2017.07.017

Dennis, N. M., Subbiah, S., Karnjanapiboonwong, A., Dennis, M. L., McCarthy, C., Salice, C. J., et al. (2021). Species- and tissue-specific avian chronic toxicity values for perfluorooctane sulfonate (PFOS) and a binary mixture of PFOS and perfluorohexane sulfonate. Environmental Toxicology and Chemistry, 40(3), 899–909. doi: 10.1002/etc.4937

Dewitt, J. C., Peden-Adams, M. M., Keller, J. M., & Germolec, D. R. (2012). Immunotoxicity of perfluorinated compounds: recent developments. Toxicologic Pathology, 40(2), 300–311. doi: 10.1177/0192623311428473

Dietz, R., Letcher, R. J., Desforges, J.-P., Eulaers, I., Sonne, C., Wilson, S., et al. (2019). Current state of knowledge on biological effects from contaminants on arctic wildlife and fish. Science of the Total Environment, 696, 1–40. doi: https://doi.org/10.1016/j.scitotenv.2019.133792

Eikenaar, C., & Hegemann, A. (2016). Migratory common blackbirds have lower innate immune function during autumn migration than resident conspecifics. Biology Letters, 12(3), 20160078. doi: 10.1098/rsbl.2016.0078

Farkas, J., Polesel, F., Kjos, M., Carvalho, P. A., Ciesielski, T., Flores-Alsina, X., et al. (2020). Monitoring and modelling of influent patterns, phase distribution and removal of 20 elements in two primary wastewater treatment plants in Norway. Science of the Total Environment, 725, 138420. doi: https://doi.org/10.1016/j.scitotenv.2020.138420

Finlayson, C. M., & Rea, N. (1999). Reasons for the loss and degradation of Australian wetlands. Wetlands Ecology and Management, 7(1), 1–11. doi: 10.1023/A:1008495619951

Fouchier, R. A. M., Bestebroer, T. M., Herfst, S., Van Der Kemp, L., Rimmelzwaan, G. F., & Osterhaus, A. D. M. E. (2000). Detection of influenza A viruses from different species by PCR amplification of conserved sequences in the matrix gene. Journal of Clinical Microbiology, 38(11), 4096–4101. doi: 10.1128/jcm.38.11.4096-4101.2000

Furness, R. W., Muirhead, S. J., & Woodburn, M. (1986). Using bird feathers to measure mercury in the environment: Relationships between mercury content and moult. Marine Pollution Bulletin, 17(1), 27–30. doi: https://doi.org/10.1016/0025-326X(86)90801-5

Gallen, C., Eaglesham, G., Drage, D., Nguyen, T. H., & Mueller, J. F. (2018). A mass estimate of perfluoroalkyl substance (PFAS) release from Australian wastewater treatment plants. Chemosphere, 208, 975–983. doi: 10.1016/j.chemosphere.2018.06.024

Garnett, S. T., Szabo, J. K., & Dutson, G. (2011). The action plan for Australian birds 2010: CSIRO Publishing.

Gebbink, W. A., & Letcher, R. J. (2012). Comparative tissue and body compartment accumulation and maternal transfer to eggs of perfluoroalkyl sulfonates and carboxylates in Great Lakes herring gulls. Environmental Pollution, 162, 40–47. doi: https://doi.org/10.1016/j.envpol.2011.10.011

Gebreeyessus, G. D., & Zewge, F. (2019). A review on environmental selenium issues. SN Applied Sciences, 1(1). doi: 10.1007/s42452-018-0032-9

Grønnestad, R., Vázquez, B. P., Arukwe, A., Jaspers, V. L. B., Jenssen, B. M., Karimi, M., et al. (2019). Levels, patterns, and biomagnification potential of perfluoroalkyl substances in a terrestrial food chain in a Nordic skiing area. Environmental Science & Technology, 53(22), 13390–13397. doi: 10.1021/acs.est.9b02533

Gu, Z., Gu, L., Eils, R., Schlesner, M., & Brors, B. (2014). circlize implements and enhances circular visualization in R. Bioinformatics, 30(19), 2811–2812.

Hansen, E., Huber, N., Bustnes, J. O., Herzke, D., Bårdsen, B.-J., Eulaers, I., et al. (2020). A novel use of the leukocyte coping capacity assay to assess the immunomodulatory effects of organohalogenated contaminants in avian wildlife. Environment International, 142, 1–8. doi: 10.1016/j.envint.2020.105861

Hargreaves, A. L., Whiteside, D. P., & Gilchrist, G. (2010). Concentrations of 17 elements, including mercury, and their relationship to fitness measures in arctic shorebirds and their eggs. Science of the Total Environment, 408(16), 3153–3161. doi: https://doi.org/10.1016/j.scitotenv.2010.03.027

Hubálek, Z. (2004). An annotated checklist of pathogenic microorganisms associated with migratory birds. Journal of Wildlife Diseases, 40(4), 639–659. doi: 10.7589/0090-3558-40.4.639

Jantawongsri, K., Nørregaard, R. D., Bach, L., Dietz, R., Sonne, C., Jørgensen, K., et al. (2021). Histopathological effects of short-term aqueous exposure to environmentally relevant concentration of lead (Pb) in shorthorn sculpin (Myoxocephalus scorpius) under laboratory conditions. Environmental Science and Pollution Research, 28(43), 61423–61440. doi: 10.1007/s11356-021-14972-6

Kelly, B. C., Ikonomou, M. G., Blair, J. D., Surridge, B., Hoover, D., Grace, R., et al. (2009). Perfluoroalkyl contaminants in an Arctic marine food web: Trophic magnification and wildlife exposure. Environmental Science & Technology, 43(11), 4037–4043. doi: 10.1021/es9003894

Kéry, M., & Schaub, M. (2011). Bayesian population analysis using WinBUGS : a hierarchical perspective (First edition. ed.): Academic Press.

Lee, J.-W., Lee, H.-K., Lim, J.-E., & Moon, H.-B. (2020). Legacy and emerging per- and polyfluoroalkyl substances (PFASs) in the coastal environment of Korea: Occurrence, spatial distribution, and bioaccumulation potential. Chemosphere, 251, 126633. doi: 10.1016/j.chemosphere.2020.126633

Liess, M., Foit, K., Knillmann, S., Schäfer, R. B., & Liess, H.-D. (2016). Predicting the synergy of multiple stress effects. Scientific Reports, 6(1), 32965. doi: 10.1038/srep32965

Lisovski, S., Gosbell, K., Minton, C., & Klaassen, M. (2020). Migration strategy as an indicator of resilience to change in two shorebird species with contrasting population trajectories. Journal of Animal Ecology, n/a(n/a). doi: https://doi.org/10.1111/1365-2656.13393

Lopez-Antia, A., Groffen, T., Lasters, R., Abdelgawad, H., Sun, J., Asard, H., et al. (2019). Perfluoroalkyl acids (PFAAs) concentrations and oxidative status in two generations of great tits inhabiting a contamination hotspot. Environmental Science & Technology, 53(3), 1617–1626. doi: 10.1021/acs.est.8b05235

Løseth, M. E., Briels, N., Eulaers, I., Nygård, T., Malarvannan, G., Poma, G., et al. (2019). Plasma concentrations of organohalogenated contaminants in white-tailed eagle nestlings – The role of age and diet. Environmental Pollution, 246, 527–534. doi: 10.1016/j.envpol.2018.12.028

Ma, Y., Choi, C.-Y., Thomas, A., & Gibson, L. (2022). Review of contaminant levels and effects in shorebirds: Knowledge gaps and conservation priorities. Ecotoxicology and Environmental Safety, 242, 113868. doi: https://doi.org/10.1016/j.ecoenv.2022.113868

Ma, Z., Melville, D. S., Liu, J., Chen, Y., Yang, H., Ren, W., et al. (2014). Rethinking China’s new great wall. Science, 346(6212), 912. doi: 10.1126/science.1257258

Marasco, V., & Costantini, D. (2016). Signaling in a polluted world: oxidative stress as an overlooked mechanism linking contaminants to animal communication. Frontiers in Ecology and Evolution, 4(95). doi: 10.3389/fevo.2016.00095

Marcelino, V. R., Wille, M., Hurt, A. C., González-Acuña, D., Klaassen, M., Schlub, T. E., et al. (2019). Meta-transcriptomics reveals a diverse antibiotic resistance gene pool in avian microbiomes. BMC Biology, 17(1). doi: 10.1186/s12915-019-0649-1

Martin, J. W., Mabury, S. A., Solomon, K. R., & Muir, D. C. G. (2003). Bioconcentration and tissue distribution of perfluorinated acids in rainbow trout (Oncorhynchus mykiss). Environmental Toxicology and Chemistry, 22(1), 196–204. doi: 10.1002/etc.5620220126

Martinez, M. P., & Kannan, K. (2018). Simultaneous analysis of seven biomarkers of oxidative damage to lipids, proteins, and DNA in urine. Environmental Science & Technology, 52(11), 6647–6655. doi: 10.1021/acs.est.8b00883

McDonough, C. A., Ward, C., Hu, Q., Vance, S., Higgins, C. P., & DeWitt, J. C. (2020). Immunotoxicity of an electrochemically fluorinated aqueous film-forming foam. Toxicological Sciences, 178(1), 104–114. doi: 10.1093/toxsci/kfaa138

Melville, D. S., Chen, Y., & Ma, Z. (2016). Shorebirds along the Yellow Sea coast of China face an uncertain future - A review of threats. Emu, 116(2), 100–110. doi: 10.1071/MU15045

Méndez, V., Alves, J. A., Gill, J. A., & Gunnarsson, T. G. (2018). Patterns and processes in shorebird survival rates: a global review. Ibis, 160(4), 723–741. doi: 10.1111/ibi.12586

Menkhorst, P., Rogers, D., Clarke, R., Davies, J., marsack, P., & Franklin, K. (2019). The Australian Bird Guide (Revised Edition): CSIRO Publishing.

Metcalfe, N. B., & Alonso-Alvarez, C. (2010). Oxidative stress as a life-history constraint: the role of reactive oxygen species in shaping phenotypes from conception to death. Functional Ecology, 24(5), 984–996. doi: 10.1111/j.1365-2435.2010.01750.x

Minton, C. (2006). The history and achievements of the Victorian Wader Study Group. Stilt, 50, 285–294.

Mitra, S., Chakraborty, A. J., Tareq, A. M., Emran, T. B., Nainu, F., Khusro, A., et al. (2022). Impact of heavy metals on the environment and human health: Novel therapeutic insights to counter the toxicity. Journal of King Saud University - Science, 34(3), 101865. doi: https://doi.org/10.1016/j.jksus.2022.101865

Murray, C. G., Kasel, S., Loyn, R. H., Hepworth, G., & Hamilton, A. J. (2013). Waterbird use of artificial wetlands in an Australian urban landscape. Hydrobiologia, 716(1), 131–146. doi: 10.1007/s10750-013-1558-x

Murray, N. J., Clemens, R. S., Phinn, S. R., Possingham, H. P., & Fuller, R. A. (2014). Tracking the rapid loss of tidal wetlands in the Yellow Sea. Frontiers in Ecology and the Environment, 12(5), 267–272. doi: 10.1890/130260

Murray, N. J., Marra, P. P., Fuller, R. A., Clemens, R. S., Dhanjal-Adams, K., Gosbell, K. B., et al. (2018). The large-scale drivers of population declines in a long-distance migratory shorebird. Ecography, 41(6), 867–876. doi: 10.1111/ecog.02957

Murray, N. J., Phinn, S. R., DeWitt, M., Ferrari, R., Johnston, R., Lyons, M. B., et al. (2019). The global distribution and trajectory of tidal flats. Nature, 565(7738), 222–225. doi: 10.1038/s41586-018-0805-8

Newsted, J. L., Jones, P. D., Coady, K., & Giesy, J. P. (2005). Avian toxicity reference values for perfluorooctane sulfonate. Environmental Science & Technology, 39(23), 9357–9362. doi: 10.1021/es050989v

Patel, M., Kumar, R., Kishor, K., Mlsna, T., Pittman, C. U., & Mohan, D. (2019). Pharmaceuticals of Emerging Concern in Aquatic Systems: Chemistry, Occurrence, Effects, and Removal Methods. Chemical Reviews, 119(6), 3510–3673. doi: 10.1021/acs.chemrev.8b00299

Picone, M., Corami, F., Gaetan, C., Basso, M., Battiston, A., Panzarin, L., et al. (2019). Accumulation of trace elements in feathers of the Kentish plover Charadrius alexandrinus. Ecotoxicology and Environmental Safety, 179, 62–70. doi: https://doi.org/10.1016/j.ecoenv.2019.04.051

Rainio, M., & Eeva, T. (2010). Metal-related oxidative stress in birds. Environmental Pollution, 158, 2359–2370. doi: 10.1016/j.envpol.2010.03.013

Rogers, K. G., Rogers, D. I., & Minton, C. D. T. (1996). Weights and pre-migratory mass gain of the red-necked stint Calidris ruficollis in Victoria, Australia. The Stilt, 29, 2–23.

Sebastiano, M., Jouanneau, W., Blévin, P., Angelier, F., Parenteau, C., Gernigon, J., et al. (2021). High levels of fluoroalkyl substances and potential disruption of thyroid hormones in three gull species from South Western France. Science of the Total Environment, 765, 1–11. doi: https://doi.org/10.1016/j.scitotenv.2020.144611

Spackman, E., Senne, D. A., Myers, T. J., Bulaga, L. L., Garber, L. P., Perdue, M. L., et al. (2002). Development of a real-time reverse transcriptase PCR assay for type A influenza virus and the avian H5 and H7 hemagglutinin subtypes. Journal of Clinical Microbiology, 40(9), 3256–3260. doi: 10.1128/jcm.40.9.3256-3260.2002

Stahl, T., Mattern, D., & Brunn, H. (2011). Toxicology of perfluorinated compounds. Environmental Sciences Europe, 23(1), 38. doi: 10.1186/2190-4715-23-38

Stark, J. S. (1998). Heavy metal pollution and macrobenthic assemblages in soft sediments in two Sydney estuaries, Australia. Marine and Freshwater Research, 49(6), 533. doi: 10.1071/mf97188

Steele, W. K., & Harrow, S. (2014). Overview of adaptive management for multiple biodiversity values at the Western treatment plant, werribee, leading to a pilot nutrient addition study. The Victorian Naturalist, 131(4), 128–146. doi: 10.3316/informit.651511656648828

Studds, C. E., Kendall, B. E., Murray, N. J., Wilson, H. B., Rogers, D. I., Clemens, R. S., et al. (2017). Rapid population decline in migratory shorebirds relying on Yellow Sea tidal mudflats as stopover sites. Nature Communications, 8(1), 1–7. doi: 10.1038/ncomms14895

Su, T., Lin, X., Huang, Q., Jiang, D., Zhang, C., Zhang, X., et al. (2020). Mercury exposure in sedentary and migratory Charadrius plovers distributed widely across China. Environmental Science and Pollution Research, 27(4), 4236–4245. doi: 10.1007/s11356-019-06873-6

Szabo, D., Marchiandi, J., Samandra, S., Johnston, J. M., Mulder, R. A., Green, M. P., et al. (2023). High-resolution temporal wastewater treatment plant investigation to understand influent mass flux of per- and polyfluoroalkyl substances (PFAS). Journal of Hazardous Materials, 447, 130854. doi: https://doi.org/10.1016/j.jhazmat.2023.130854

Trimmel, S., Vike-Jonas, K., Gonzalez, S. V., Ciesielski, T. M., Lindstrøm, U., Jenssen, B. M., et al. (2021). Rapid determination of per- and polyfluoroalkyl substances (PFAS) in harbour porpoise liver tissue by HybridSPE®–UPLC®–MS/MS. Toxics, 9(8), 1–11. doi: 10.3390/toxics9080183

Tsipoura, N., Burger, J., Niles, L., Dey, A., Gochfeld, M., Peck, M., et al. (2017). Metal levels in shorebird feathers and blood during migration through Delaware Bay. Archives of Environmental Contamination and Toxicology(4), 562. doi: 10.1007/s00244-017-0400-2

UNEP. (2022). Stockholm Convention on persistent organic pollutants. Retrieved 18 August 2022, 2022, from http://chm.pops.int

US Environmental Protection Agency. (2022). PFAS structures in DSSTox (update August 2022).

Valavanidis, A., Vlahogianni, T., Dassenakis, M., & Scoullos, M. (2006). Molecular biomarkers of oxidative stress in aquatic organisms in relation to toxic environmental pollutants. Ecotoxicology and Environmental Safety, 64(2), 178–189. doi: https://doi.org/10.1016/j.ecoenv.2005.03.013

Vallverdú-Coll, N., Mateo, R., Mougeot, F., & Ortiz-Santaliestra, M. E. (2019). Immunotoxic effects of lead on birds. Science of the Total Environment, 689, 505–515. doi: https://doi.org/10.1016/j.scitotenv.2019.06.251

Verhagen, J. H., Herfst, S., & Fouchier, R. A. M. (2015). How a virus travels the world. Science, 347(6222), 616–617. doi: doi:10.1126/science.aaa6724

Wang, Z., Dewitt, J. C., Higgins, C. P., & Cousins, I. T. (2017). A never-ending story of per- and polyfluoroalkyl substances (PFASs)? Environmental Science & Technology, 51(5), 2508–2518. doi: 10.1021/acs.est.6b04806

Warnock, S. E., & Takekawa, J. Y. (2008). Wintering site fidelity and movement patterns of Western Sandpipers Calidris mauri in the San Francisco Bay estuary. Ibis, 138(2), 160–167. doi: 10.1111/j.1474-919x.1996.tb04323.x

Wille, M., Eden, J.-S., Shi, M., Klaassen, M., Hurt, A. C., & Holmes, E. C. (2018). Virus–virus interactions and host ecology are associated with RNA virome structure in wild birds. Molecular Ecology, 27(24), 5263–5278. doi: 10.1111/mec.14918

Wille, M., Grillo, V., Ban De Gouvea Pedroso, S., Burgess, G. W., Crawley, A., Dickason, C., et al. (2021). Australia as a global sink for the genetic diversity of avian influenza A virus. Cold Spring Harbor Laboratory. Retrieved from https://dx.doi.org/10.1101/2021.11.30.470533

Wille, M., Lisovski, S., Risely, A., Ferenczi, M., Roshier, D., Wong, F. Y. K., et al. (2019). Serologic evidence of exposure to highly pathogenic avian influenza H5 viruses in migratory shorebirds, Australia. Emerging Infectious Diseases, 25(10), 1903–1910. doi: 10.3201/eid2510.190699

Wille, M., Lisovski, S., Roshier, D., Ferenczi, M., Hoye, B. J., Leen, T., et al. (2023). Strong host phylogenetic and ecological effects on host competency for avian influenza in Australian wild birds. Proceedings of the Royal Society B: Biological Sciences, 290(1991), 20222237. doi: doi:10.1098/rspb.2022.2237

Winkel, L. H. E., Johnson, C. A., Lenz, M., Grundl, T., Leupin, O. X., Amini, M., et al. (2012). Environmental selenium research: from microscopic processes to global understanding. Environmental Science & Technology, 46(2), 571–579. doi: 10.1021/es203434d

Yue, S., Huang, C., Wang, R., & Qiao, Y. (2021). Selenium toxicity, bioaccumulation, and distribution in earthworms (Eisenia fetida) exposed to different substrates. Ecotoxicology and Environmental Safety, 217, 112250. doi: https://doi.org/10.1016/j.ecoenv.2021.112250

Zhang, J., Sundfør, E. B., Klokkerengen, R., Gonzalez, S. V., Mota, V. C., Lazado, C. C., et al. (2022). Determination of the oxidative stress biomarkers of 8-hydroxydeoxyguanosine and dityrosine in the gills, skin, dorsal fin, and liver tissue of atlantic salmon (Salmo salar) parr. Toxics, 10(9), 1–13. doi: 10.3390/toxics10090509

Zhang, Q., & Wang, C. (2020). Natural and human factors affect the distribution of soil heavy metal pollution: A review. Water, Air, & Soil Pollution, 231(7), 1–13. doi: 10.1007/s11270-020-04728-2

